# Purification, identification, and characterization of two benzophenanthridine alkaloids from *Corydalis longicalcarata* rhizomes with anti-mitotic and polyploidy-inducing activities

**DOI:** 10.1101/821124

**Authors:** Jinhua Li, Ziqi Yan, Hongmei Li, Qiong Shi, Linfang Huang, Thaddeus D. Allen, Dun Yang, Jing Zhang

**Affiliations:** J. Michael Bishop Institute of Cancer Research, Chengdu, China; Institute of Medicinal Plant Development, Peking Union Medical College & Chinese Academy of Medical Sciences, Beijing, China

## Abstract

Mitosis has long been a therapeutic target for the treatment of cancer. After conducting a phenotypic screen for anti-mitotic activity in a crude extract library of more than 2000 medicinal plants, we identified the rhizomes of *Corydalis longicalcarata*. Guided by the bioactivity-based assay, two benzophenanthridine alkaloids were purified and subsequently identified as corynoline and its close analog acetylcorynoline. This study uncovers a previously unknown antimitotic activity for these two phytochemicals, discovers their potential to be developed as anticancer drugs, and highlights their pleiotropic effects on cell division, including prevention of chromosome congression, compromise of spindle checkpoint response, and blockage of cytokinesis.

## INTRODUCTION

The mitotic spindle is the dynamic microtubule structure that segregates chromosomes during mitosis. The mitotic structure has long been a therapeutic target for the treatment of cancer and spindle toxins such as taxanes and vinca alkaloids have been used clinically for decades. Taxanes work by stabilizing microtubules and halting depolymerization whereas vinca alkaloids function to inhibit tubulin polymerization. Despite their distinct mechanisms of action, both types of spindle toxins interfere with the function of the mitotic spindle, leading to the persistent activation of the spindle checkpoint response. This, in turn, causes a prolonged arrest of cells in mitosis and eventually cell death by apoptosis or mitotic catastrophe. Despite their success as chemotherapeutics, microtubule-interfering drugs exert the mechanism-based cytotoxicity to the central nervous system when used at therapeutic doses. An alternative approach that is potentially less toxic to target mitotic regulators of the spindle assembly, rather than the spindle microtubules *per se*. This may achieve similar therapeutic efficacy but overcome the adverse effects associated with spindle toxins.

We conducted a screening for compounds with novel anti-mitotic activity in a library of crude herbal extracts. Specifically, our screen was focused on identifying extracts that could activate the spindle checkpoint and arrest cells in mitosis. We identified an ethanol extract from the rhizomes of *Corydalis longicalcarata* (Papaveraceae) with the desired activity. In the present work, we conducted bioassay-guided isolation and purification of this anti-mitotic activity. The bioactive molecules were identified as corynoline and acetylcorynoline. Both molecules have been previously isolated from *Corydalis incisa* ^1^ and *Corydalis bungeana* ^2^ and are known to have a variety of pharmacological activities. Known activities include acetylcholinesterase inhibitory activity ^3^, anti-fungal activity ^4^, inhibition of cell adhesion ^5^, anti-inflammatory properties^6^, ability to inhibit β-site amyloid precursor protein cleaving enzyme 1 (BACE1) ^7^ and cytotoxic activities ^8^. However, to our knowledge, an anti-mitotic activity for these two phytochemicals has not been reported. We found that both phytochemicals elevate the mitotic index, transiently arrest cells in prometaphase, and ultimately induce polyploidy by preventing cytokinesis. For each activity, corynoline has a minimal effective concentration 2-fold lower than that of acetylcorynoline. The anti-mitotic effects of these phytochemicals are distinct from those of spindle toxins. Cells exposed to spindle toxins undergo a prolonged arrest in mitosis. Our findings indicate that corynoline and acetylcorynoline interfere with mitosis by a mechanism that is distinct from that utilized by spindle toxins and that these molecules can serve as prototypic molecules with the potential to be developed as more potent and specific anticancer therapeutics.

## RESULTS AND DISCUSSION

We have constructed a natural products library composed of 17,000 crude extracts from more than 2,000 different plant species. Using this library, we conducted a phenotypical screen for anti-mitotic activity. Cellular “round-up” was used as a surrogate marker for mitotic arrest and could easily be assayed using an inverted tissue culture microscope (Figure S1). One extract, made from the rhizomes of *C. longicalcarata* (#1779) was identified as positive, retested, and confirmed to possess multiple anti-mitotic activities in subsequent assays. These included initial provocation of the spindle checkpoint response, followed by compromising its maintenance and ultimately induction of polyploidy. In this study, we purified, identified, and characterized the bioactive ingredients from the *C longicalcarata* rhizome that exerts these pleiotropic effects on cell division.

The success of our fractionation and eventual identification of bioactive components was aided by the pre-study design of a simple 96-well plate screening assay. As cells enter and become arrested in mitosis, they detach from the plate surface and appear as rounded cells with smooth surface membrane, distinct from detached cells that are undergoing programmed cell death. We utilized this simple visual screening method to assay thousands of extracts. It should be noted that although activity was assayed in other parts of *C. longicalcarata*, our rhizome extract was found to contain substantial activity. This may indicate a preponderance for this portion of the plant to synthesize and/or store the bioactive compounds compared to other portions of the plant.

To identify bioactive components that are able to interfere with mitosis, we recollected 3 kg of *C. longicalcarata* rhizome and performed bioactivity-guided extraction, isolation, and purification (Figure S2). The air dried and powdered material was extracted with 70% ethanol assisted by exposure to ultrasonic waves. This liquid was then partitioned sequentially with petroleum ether (PE) and ethyl acetate (EA) to yield, in total, 55 distinct fractions by MPLC. These were again assayed for anti-mitotic activity and we found 10 PE and 3 EA fractions with activity (Figure S3). Some fractions were active at lower concentrations, but not at higher. This was deemed due to impurities that at were able to alter our assay outcome and we sought to further purify the bioactive fractions using TLC (Figure S4). Of the 62 PE and 22 EA TLC fractions, we found 11 retained mitotic activity in our 96-well plate assay at the lowest concentration assayed, 6.25 μg / ml (Figure S5).

As a next step we performed analytical HPLC with the 8 positive PE sub-fractions and 3 positive EA sub-fractions, revealing that 7 sub-factions in total possessed a shared peak (we have designated peak 1), whereas the remaining four sub-fractions a shared a second peak (we have designated peak 2) (Figure S6). The presence of two shared peaks among the 11 sub-fractions indicated that at least two natural compounds were responsible for the antimitotic activity in our biological screening assays. The peak 1-containing sub-fractions were pooled and loaded onto a preparative HPLC to obtain a natural compound 1 with 88% purity (Figure S7). Likewise, compound 2 with a purity of 90% was purified from the pooled sub-fractions that contain peak 2 (Figure S7).

We performed physiochemical analysis of compounds 1 and 2, and then asked if their chemical profiles match those of any known compounds isolated from plants of the genus *Corydalis* in the literature. Although compounds 1 and 2 had not matched any phytochemicals reported in *C. longicalcarata*, they matched corynoline and acetylcorynoline, respectively, which have been identified from two other species of the genus *Corydalis, Corydalis incisa* ^1^ and *Corydalis bungeana* ^2^. Three lines of evidence supported such identification. First, as determined by the mass spectrometry, the molecular weight of compounds 1 and 2 are 367 and 409, respectively, matching those of corynoline and acetylcorynoline (Figure S8). Second, the two compounds are also indistinguishable from corynoline and acetylcorynoline in ^1^H NMR,^13^C NMR and UV spectra (Table S1 and data not shown). Third, analytic HPLC analysis demonstrated that compound 1 and the commercially obtained corynoline eluted at the same time of 27.07 min, whereas compound 2 and the commercially obtained acetylcorynoline had the exact the same elution time of 37.59 min (Figure S9).

Plants produce various secondary metabolites that fall into major phytochemical classes including flavonoids, tannins, terpenoids, saponins, triterpenoid saponins, alkaloids, phytosterols, carotenoids, fatty acids and essential oils <u>^11^</u>. The main classes of metabolites that have been transformed into modern drugs include terpenes (34%), glycosides (32%), alkaloids (16%) and others (18%). Corynoline and acetylcorynoline (Figure S10) fall into the category of benzophenanthridine alkaloids, and have been reported to have various pharmacological properties. But whether they have antimitotic activity has not been reported.

We confirmed their effect on cell-division using corynoline and acetylcorynoline purified either in-house or commercially. Cells treated with either of these compounds suffered similar cell-division defects. Nearly all cells entered and halted in mitosis 24 hours after the initiation of treatment (Figure S11a-d). The majority of the cells had condensed DNA without apparent alignment of chromosomes on the metaphase plate, indicating that the mitotic spindle checkpoint was activated and subsequently arrest was in prometaphase. The arrest of cells in mitosis was, however, transient. Polyploidy prevailed 48 hours after the initiation of the treatment (Figure S11e and Figure S12). Thus, the mitotic spindle checkpoint was activated but failed to be maintained in the presence of corynoline or acetylcorynoline. Treated cells eventually exited from mitosis without completion of cytokinesis, leading to the formation of multinucleated cells.

Threshold concentrations of corynoline to elicit mitotic arrest and polyploidy were indistinguishable, indicating that disablement of the same mitotic regulator might be responsible for both effects (Table S2). Acetylcorynoline mimicked corynoline in eliciting both mitotic arrest and polyploidy (Figure S13), although the minimal effective concentration of acetylcorynoline was a 2-fold higher than that of corynoline (Table S2). Aurora-B kinase, the catalytic subunit of the chromosomal passenger protein complex, plays an essential role in coordinating chromosome segregation with cytokinesis <u>^12^</u>. However, the effect of corynoline on cell division was not mediated by inhibition of aurora-B kinase because the phosphorylation of Histone H3 at Ser10, a surrogate of aurora-B kinase activity ^13, 14^ was not affected by either corynoline or acetycorynoline (Figure S11b and Figure S12A). In contrast with corynoline and acetylcorynoline, both taxol and vinblastine elicited a persistent arrest of cells in mitosis without causing accumulation of polyploid cells within 48-hour treatment (Figure S13 and Table S3). Thus, corynoline and acetylcorynoline warrant further study as antimitotic therapeutics for the treatment of cancer. They possess a unique anti-mitotic activity that may be advantageous to harness as a chemotherapeutic.

The chemical composition and associated medicinal properties of *C. longicalcarata* are uncharacterized. Using a bioactivity-guided assay, we isolated and purified, two antimitotic phytochemicals from the rhizomes of *C. longicalcarata* and identified them as corynoline and its derivative acetylcorynoline. Although these two phytochemicals have been previously identified from other species of the genus *Corydalis*, this is the first report that they are found in *C. longicalcarata* and have a previously unknown antimitotic activity. The phytochemicals both elicit pleiotropic effects on cell division including prevention of chromosome segregation, compromising the spindle checkpoint response, and blockade of cytokinesis. These cell-division effects warrant further study of these compounds as anticancer therapeutics.

## EXPERIMENTAL SECTION

### Plant materials

The rhizomes of *Corydalis longicalcarata* were collected in April 2018 from Longchi National Forest Park, located in the northwest of Dujiangyan, Sichuan Province, China (altitude: 1625 m; latitude: N 31°6’10”; longitude: E 103°33’20”). This original plant was authenticated as *Corydalis longicalcarata* by professor Linfang Huang, Institute of Medicinal Plant Development, Peking Union Medical College & Chinese Academy of Medical Sciences, Beijing, China. A voucher specimen was assigned a number of MBICR-0728 and deposited at the Herbarium of the J. Michael Bishop Institute of Cancer Research (MBICR).

### Chemicals and reagents

The reference standards for corynoline and acetylcorynoline were purchased from Chengdu Must Biotechnology Co. Ltd. (Sichuan, China). The purity of these two natural compounds were higher than 98% as validated by analytic HPLC in-house. Silica gel (200-300 mesh) and all solvents used for extraction and isolation were from Chengdu Kelong Chemical Co., Ltd (Sichuan, China). Preparative TLC plates (silica gel, 200 mm × 200 mm × 1mm) were obtained from Yantai Xinnuo Co. Ltd. (Shandong, China). All HPLC grade solvents were from Fisher Scientific (Fisher Scientific, USA) and used without further purification. Taxol and Vineblastine were purchased from Sigma. De-ionized water was purified using a Milli-Q system (Millipore, Bedford, MA, USA).

### Cell culture and treatment

Retinal pigment epithelial cells transformed by ectopic MYC and Bcl-2 expression (RPE-MBC cells) have been described previously ^9^ and were cultured in DMEM supplemented with 5% fetal bovine serum in a humidified incubator that was maintained at 5 % CO_2_. For screening for bioactive fractions, cells were cultured in 96-well plates, and exposed to crude extracts, partially purified fractions or pure compounds at the concentrations indicated in the figure legends. Exposure was for 24 or 48 hours before analysis of cells for either an arrest in mitosis using an inverted microscope or for a change in DNA content after 4’,6’-diamidino-2-phenylindole (DAPI) staining by fluorescence microscopy.

### Immunofluorescent staining

Immunofluorescence staining was with a mouse monoclonal antibody against Histone H3 phosphorylated at Ser10 and has been previously described <u>^10^</u>. Cells were cultured on coverslips in a 6-well plate, fixed with 4 % paraformaldehyde and then permeabilized with 0.5 % Triton X-100 in PBS. Primary antibody was detected with Texas-Red-conjugated secondary antibodies that were purchased from Jackson ImmunoResearch. After immunostaining, we mounted cells on microscope slides with 4’,6’-diamidino-2-phenylindole (DAPI)-containing Vectashield mounting solution (Vector Laboratories) for detection of fluorescence with an EVOS FL fluorescence microscope (ThermoFisher).

### Extraction, isolation and purification of compounds 1 and 2

Samples from *C. longicalcarata* were dried in air, chopped, and ground to fine powder in an electric grinder. Three kilograms (kg) of powder was extracted with 70% ethanol under ultrasound treatment (40 KHz) for 0.5 hours at room temperature, and then without ultrasound treatment for another 24 hours. The extraction procedure was repeated 3 times. Each time, the mixture was filtered, solvent was evaporated under vacuum using a rotary evaporator (N-1300, Tokoyo Rikakikai Co. Ltd), and the residue was dissolved in deionized water before successively partitioned against petroleum ether (PE) and ethyl acetate (EA) to obtain PE (PE, 8.5 g) and EA (EA, 9.7 g) phases, respectively.

After evaporation under vacuum, PE and EA phases were separated by column chromatograph to obtain 30 (PE1-30) and 25 (EA1-25) distinct fractions, respectively. During this process, the PE phase was loaded onto a silica gel column (100 g) and eluted manually with a step-wise gradients of petroleum ether - ethyl acetate solution (10:1, 5:1, 3:1, 2:1, 1:1, 1:2, 1:3, 1:4, 0:1, v/v) to generate the 30 fractions (PE1-30). The EA phase was subjected to low pressure preparative liquid chromatograph (SepaBean™ Machine, Santai Technologies Inc.) on a silica gel flash column (330 g). Elution was at a flow rate of 60 ml/min to produce the 25 fractions (EA1-25) with the following petroleum ether (A) - ethyl acetate (B) gradient: 0-5 min, 0→5 % B; 5-20 min, 5→30 % B; 20-30 min, 30→50 % B; 30-60 min, 50→80 % B; 60-90 min, 80→100 % B.

Next, the anti-mitotic activity present in each fraction was monitored by *in vitro* assay. Ten fractions from the PE phase and three fractions from the EA phase were positive in our assays at a concentration of 12.5 μg / ml and these fractions were further separated using preparative silica gel TLC plates to yield 62 PE and 22 EA subfractions of further purity. TLC analyses that were carried out on preparative TLC plates (silica gel, 200 mm × 200 mm × 1mm) (Yantai Xinnuo Co. Ltd.) were examined with UV light at 254 and 365 nm, followed by spraying with the Dragendorff’s reagent.

Eight out of the 62 PE subfractions and 3 out of the 22 EA sub-fractions tested positive for arresting cells in mitosis at a concentration of 6.25 μg / ml and fell into either of two groups based on whether their analytic HPLC spectra contain a shared peak, designated as peaks number 1 or 2 (HPLC: 1260 Infinity II LC System (Agilent); the column: Waters Xbridge C18, 4.6 mm × 250 mm, 5 µm (Waters); mobile phase: A, 0.2 % acetic acid-triethylamine solution (pH 5.0); B, acetonitrile. 0-5 min, 10 % B; 5-10 min, 10→20 % B; 10-70 min, 20→60 % B; 70-75 min, 60→10 % B; 75-90 min, 10 % B; column temperature: 30 °C; flow rate: 1 ml / min; the injection volume: 10 *µ*l).

Further preparative HPLC (LC-20AP, Shimadzu Corp.) for separation of these active sub-fractions was conducted on a Shimadzu Shim-pack PRC-ODS C18 column, 20 mm × 250 mm, 5 µm (mobile phase: A, 0.2 % acetic acid-triethylamine solution (pH 5.0); B, acetonitrile. 0-5 min, 10 % B; 5-10 min, 10→20 % B; 10-70 min, 20→60 % B; 70-75 min, 60→10 % B; 75-90 min, 10 % B; column temperature: 30 °C; flow rate: 18 ml / min) and this enabled purification of compound 1 (18 mg) and compound 2 (21 mg), corresponding to the peaks 1 and 2, respectively.

### Physiochemical properties of compounds 1 and 2

The structures of compounds obtained were determined by the analysis of ^1^H and ^13^C NMR spectra, ESI MS and UV absorbency spectrum. All ^1^H-NMR and ^13^C-NMR spectra were recorded in CDCl_3_ on a Bruker Avance III 400 MHz NMR operating at 400 MHz for ^1^H and 100 MHz for ^13^C. Chemical shifts were recorded as δ values in parts per million (ppm) and were normalized to tetramethylsilane (0.00 ppm) as an internal standard. Chemical shift multiplicities are reported as s = singlet, d = doublet, t = triplet, q = quartet and m = multiplet. Coupling constants (J) are given in Hz.

The ^13^C NMR spectra revealed 21 and 23 carbon atoms in compounds 1 and 2, respectively. ^1^H NMR spectrum of compound 2 is quite similar to that of compound 1. The specific chemical shifts in the ^1^H NMR at δ 1.15 (3H, s) for compound 1 and δ 1.27 (3H, s) for compound 2 imply the existence of a tertiary methyl in both compounds. Furthermore, both compounds 1 and 2 show an N-methyl singlet at δ 2.23 (3H), two doublets for two methylenedioxy groups at δ 5.97, 6.00 (4H), two aromatic singlets at δ 6.65, 6.66 (2H), and a typical AB quartet at δ 6.79, 6.91 (2H, J=8.3 Hz). Compound 2 differs from compound 1 in having additional chemical shifts in ^1^H NMR at δ 1.86 (3H, s) and ^13^C NMR at δ 169.7 and 20.8, indicating the existence of an - OCOCH_3_ group.

LC-MS analyses were performed using the Agilent 6120 Quadrupole LC/MS instrument (Agilent Technologies). Waters Xbridge C18 columns (4.6 mm × 50 mm, 5 µm) were used. Mobile phase: A, water (0.01 mol/L NH_4_HCO_3_); B, acetonitrile. Gradient: B from 5% to 100% for 1.6 min and hold 100% for 1.4 min. Column temperature: 40 °C; flow rate: 2 ml/min; the injection volume: 1 µL. NMR data together with their molecular weight of 367 and 409 revealed by MS spectrum (m/z 368.2 [M+H]^+^ and 410.2 [M+H]^+^) lead to determination of molecular formulas for compounds 1 and 2 as C_21_H_21_NO_5_ and C_23_H_23_NO_6_, respectively.

Characteristic UV absorption spectra were acquired on a Nanodrop (Thermo Fisher) and showed maximum absorption peaks at 290 nm for both compounds dissolved in methanol.

Comparison of these data with published data ^4, 6^ and the Pubchem database led to the conclusion that compounds 1 and 2 are corynoline and acetylcorynoline, respectively. We further confirmed the conclusion by comparative analysis of compounds 1, 2 and commercially available corynoline and acetylcorynoline with analytic HPLC. The purity verified by HPLC / NMR was greater than 87 % for the purified compounds 1 and 2 and 97 % for commercial obtained corynoline and acetylcorynoline.

## ASSOCIATED CONTENT

### Supporting Information

Supplemental Figures (.docx)

Supplemental Tables (.docx)

## AUTHOR INFORMATION

### Corresponding Author

Correspondence should be addressed to.

### Author Contributions

The manuscript was written through contributions of all authors. All authors have given approval to the final version of the manuscript.

### Funding Sources

This work is supported by an endowment fund to MBICR.

## ACKNOWLEDGEMENT

The authors would like to thank Yan Long for technical support and the chemistry group at MBICR for discussion.

## ABBREVIATIONS

DMSO: Dimethyl Sulfoxide
EtOH: ethanol
HPLC: high pressure liquid chromatography
LC-MS: liquid chromatography mass spectrometry
MeOH: methanol
NMR: nuclear magnetic resonance spectroscopy
TLC: thin layer chromatography.
MBICR: J. Michael Bishop Institute of Cancer Research.

## Visual Abstract

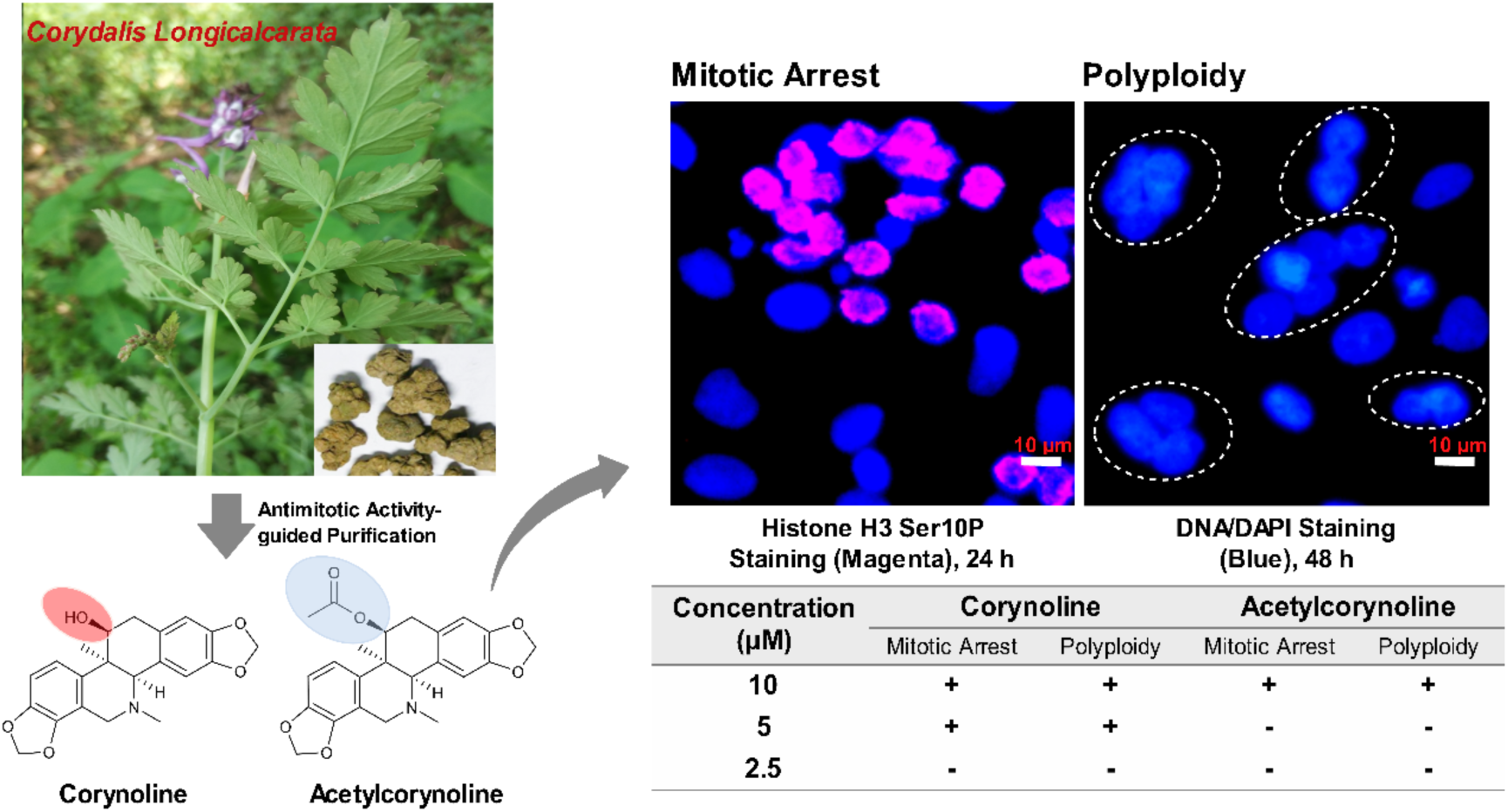

**Figure S1.**
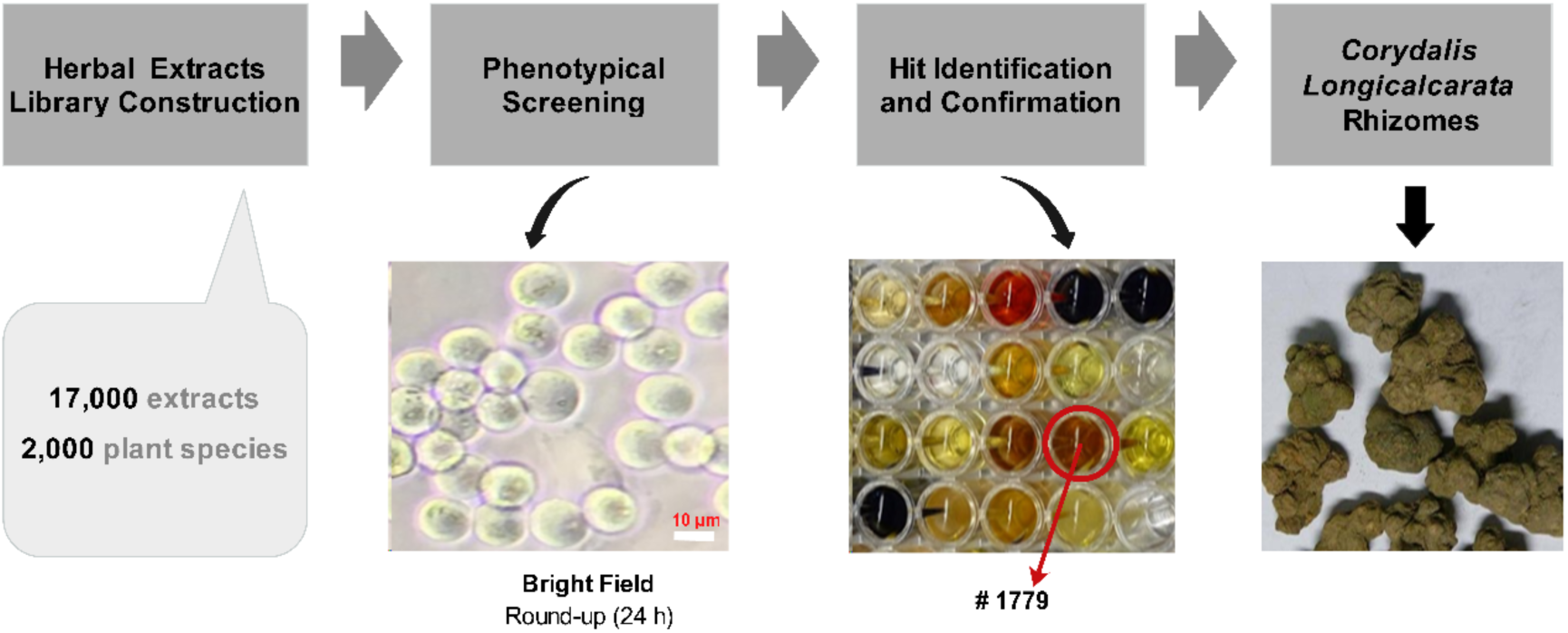
A Phenotypic screen of a natural products library identified an antimitotic activity in *C. longicalcarata* rhizomes. A crude extract library of 17,000 samples from 2,000 plant species was phenotypically screened in 96-well microplates seeded with transformed RPE-MBC cells for ability to arrest cells in mitosis. Mitotic arrest was judged by cell round-up observed under an inverted tissue culture microscope 24 hours after exposing cells to the library. One active extract made from the rhizomes of *C. longicalcarata* (#1779) was first identified, then re-assayed and activity confirmed.

**Figure S2.**
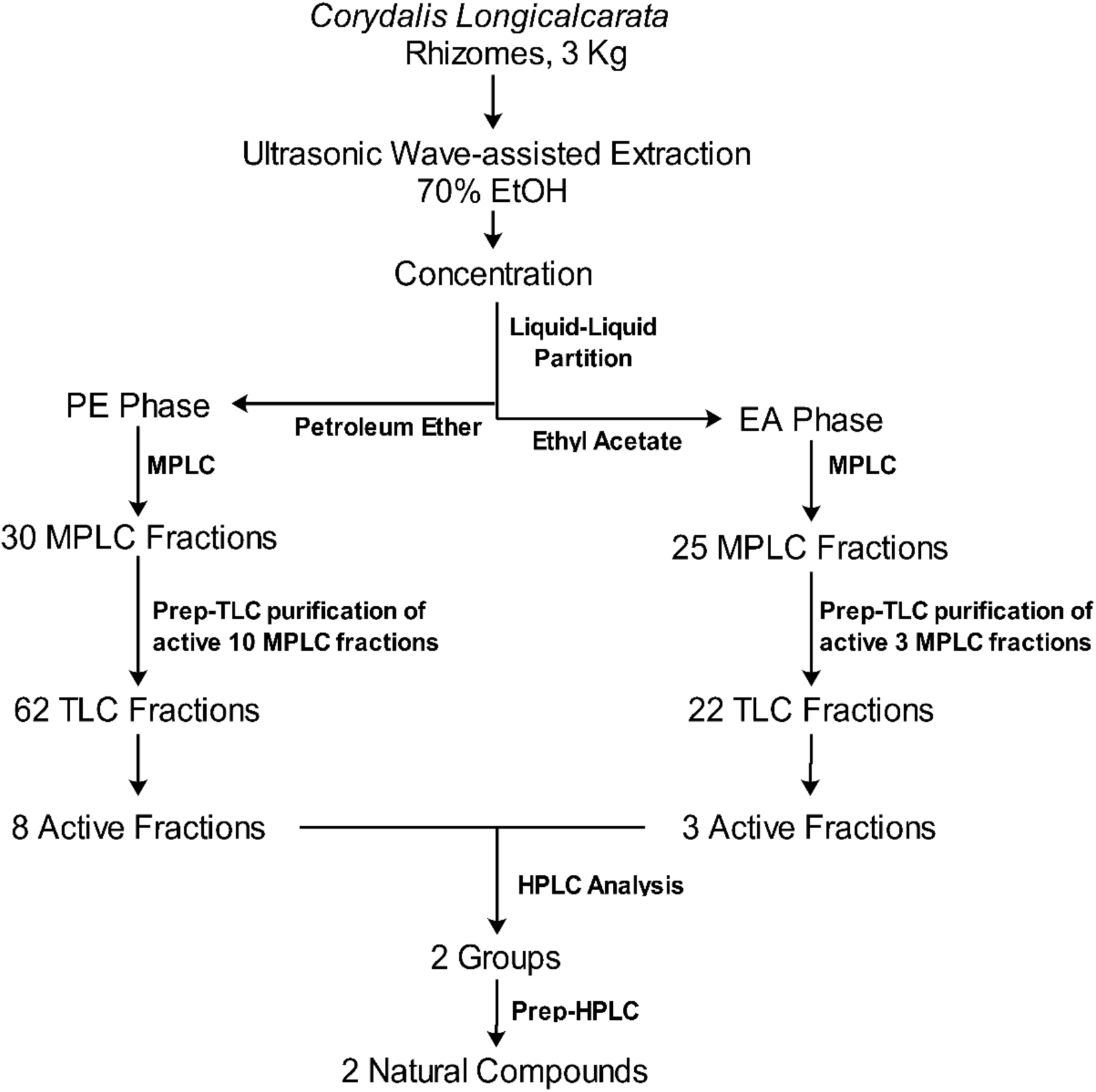
Flow chart describing steps used in the isolation and purification of natural compounds with mitotic arrest / polyploidy-inducing activity from *C. longicalcarata*.

**Figure S3.**
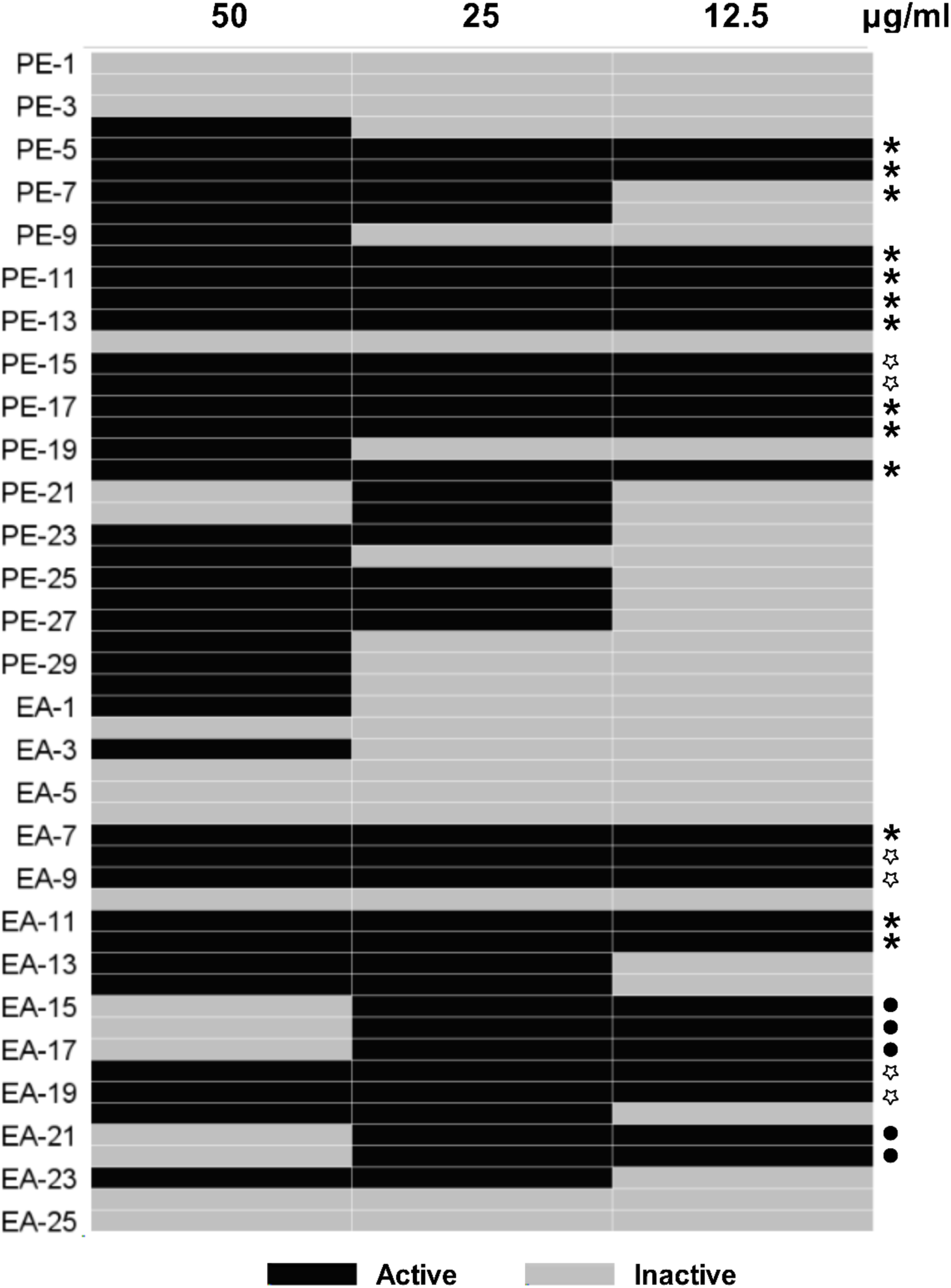
The antimitotic activity of partially purified MPLC fractions in RPE-MBC cells. Cells were exposed to MPLC fractions (30 PE fractions and 25 EA fractions) at the indicated concentrations for 24 hours before analysis for mitotic arrest using cell round-up as a surrogate. Active and inactive fractions were defined as those that elicited cell round-up in more than or less than 30% of population, respectively. *, Fractions subjected to further purification by Prep-TLC. •, Fractions positive at low concentrations but not at the highest concentration., Fractions have potent antimitotic activities but their bioactive phytochemicals are yet to be purified.

**Figure S4.**
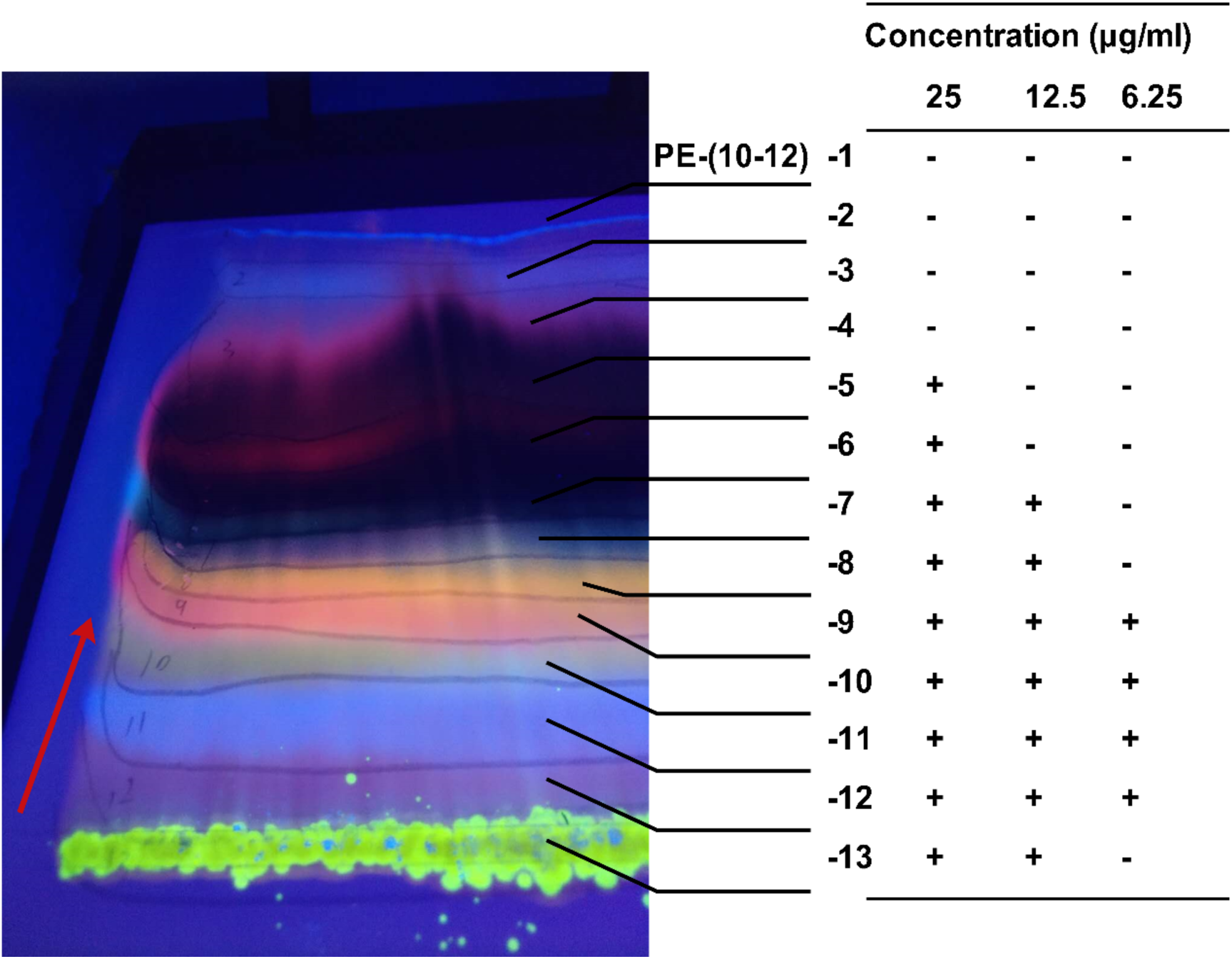
Purification of the anti-mitotic activity using preparative thin layer chromatography (prep-TLC). A representative TLC is shown in which MPLC fraction, PE-(10-12) (311 mg), was loaded on a preparative TLC plate and separated using PE-EA (1:1) as a running solution. Thirteen fractions with distinct fluorescence emission wavelength were observed under UV excitation of 365 nm and photographed. The red arrow indicates the migration direction of running solution. Each of these fractions was cut out and incubated with methanol to recover phytochemicals for the determination of their activities in eliciting mitotic arrest (activity indicated at right). The symbols + and – indicate with or without ability to cause cells to round-up, respectively.

**Figure S5.**
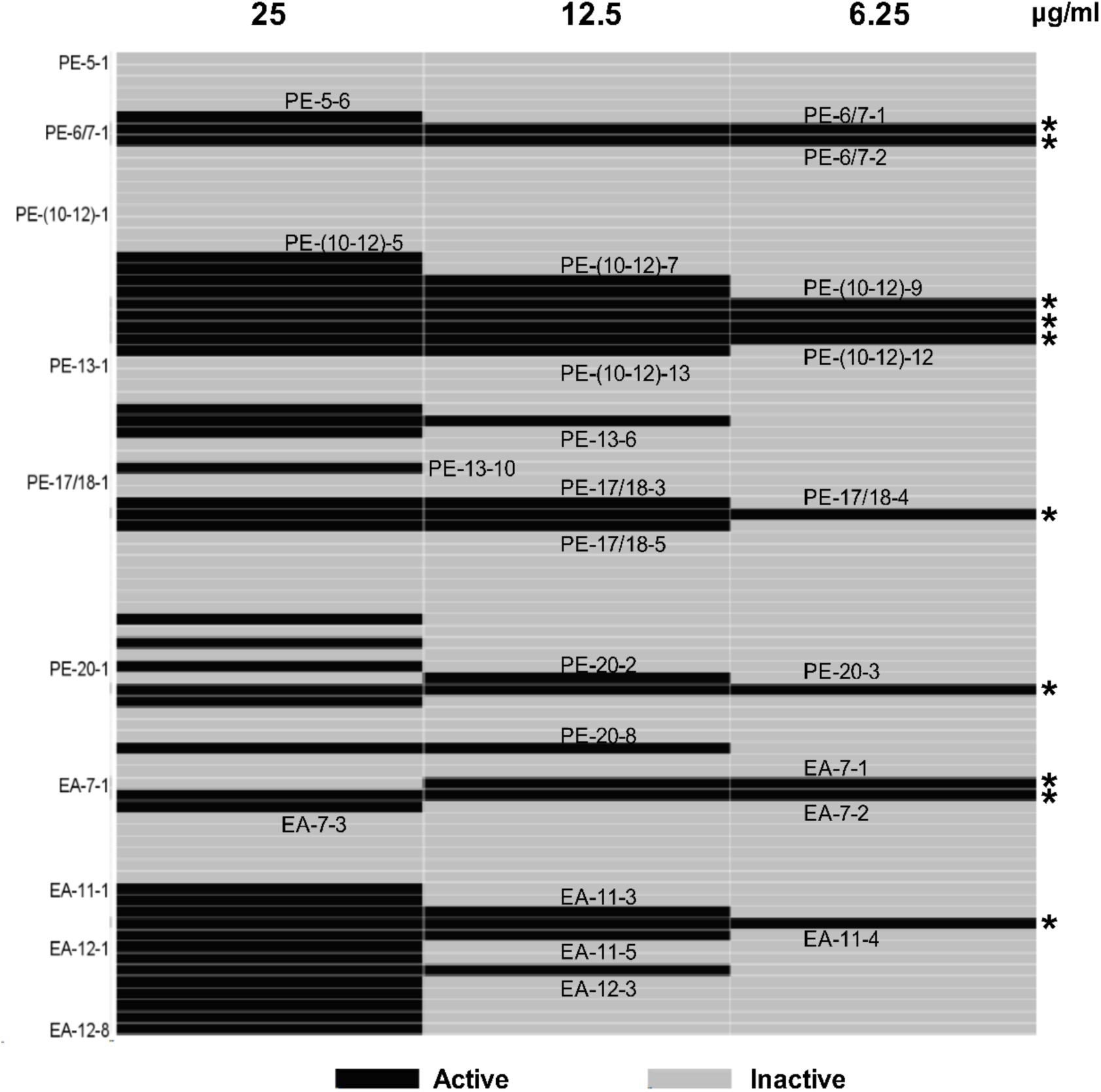
The antimitotic activity of TLC fractions. The 62 new prep-TLC fractions derived from the 10 active petroleum ether phase MPLC fractions and the 22 new prep-TLC fractions derived from the 3 active ethyl acetate phase MPLC factions, were assayed anew for their antimitotic activity. RPE-MBC were exposed to the 84 prep-TLC fractions at the indicated concentrations for 24 hours before observation of cell round-up under an inverted microscope. Active and inactive fractions are defined as those that elicit cell round-up in more than or less than 30% of population, respectively. *, Fractions with the highest antimitotic activity.

**Figure S6.**
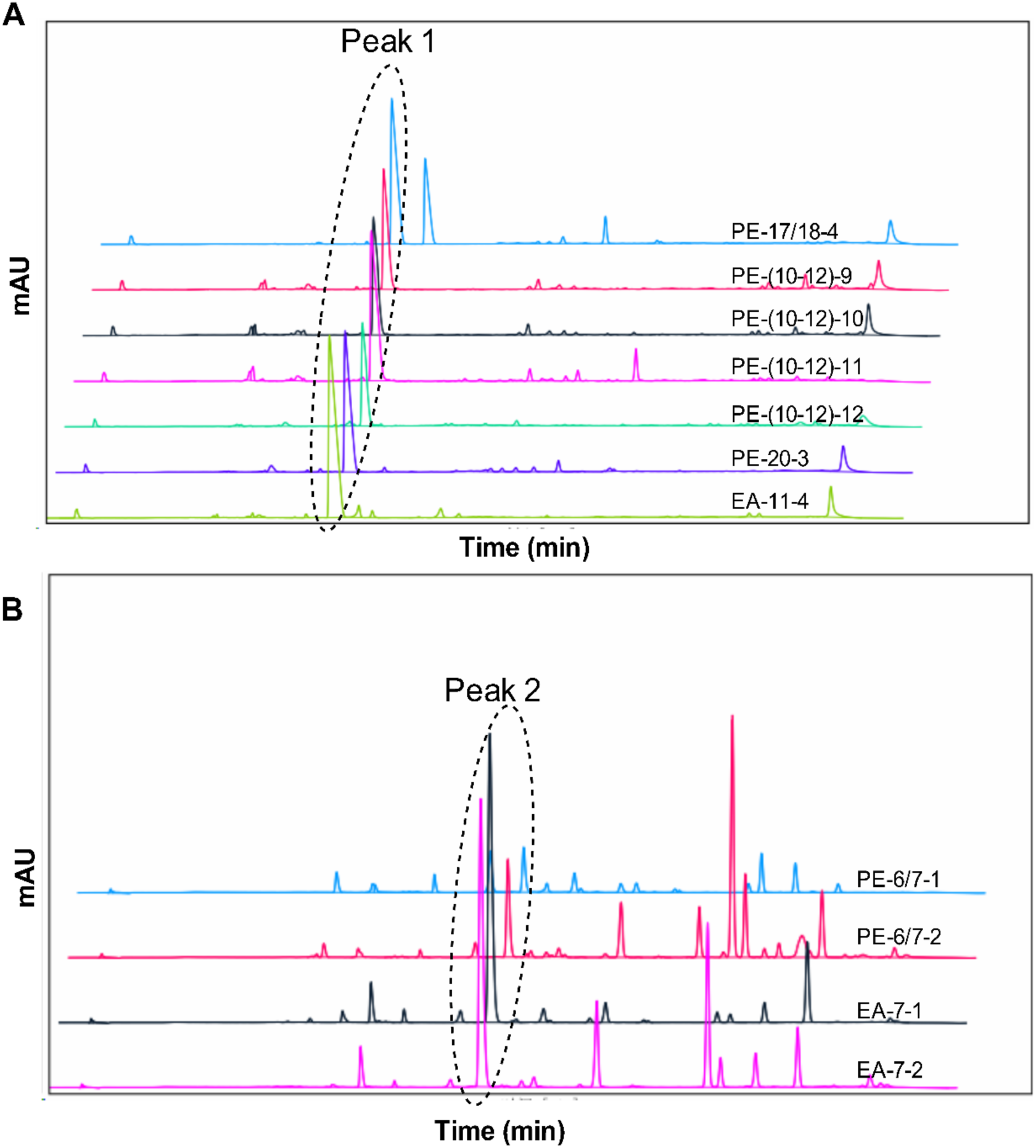
Analytical HPLC analysis of the 11 active fractions from prep-TLC. Samples were analyzed by HPLC at 280 nm. Seven fractions exhibit a shared peak 1 (A) and the remaining 4 fractions a shared peak 2 (B).

**Figure S7.**
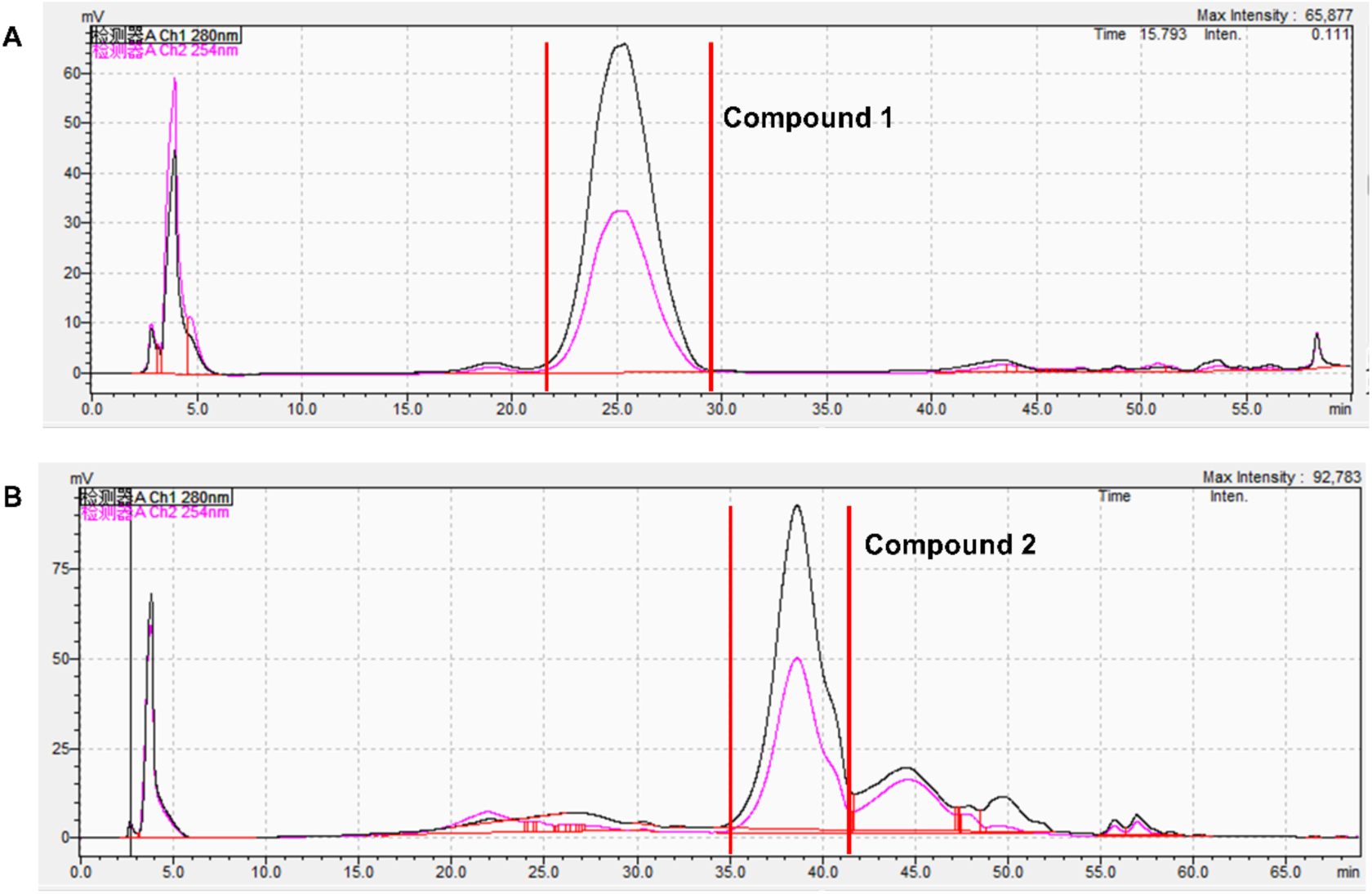
Purification of compounds 1 and 2 by preparative HPLC. Peak 1-containing fractions were combined and loaded onto a preparative HPLC column to purify compounds 1 (A). Likewise, peak 2-containing fractions were used to purify compounds 2 (B).

**Figure S8.**
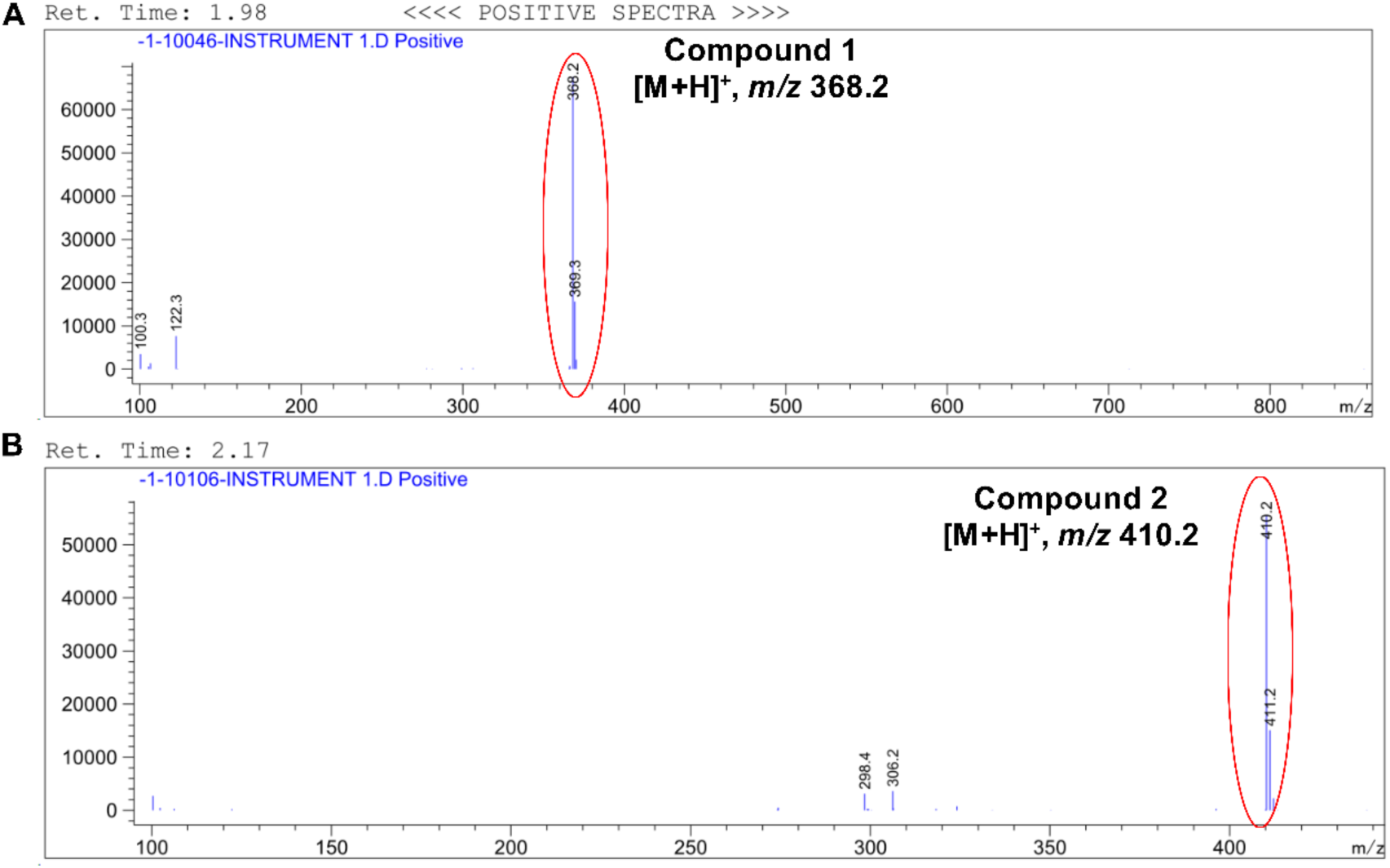
MS spectra of the isolated compound 1 (A) and compound 2 (B). Compound 1 gave an estimated MW of 368.2 and compound 2 a MW of 410.2, very close to 367 and 409, the MWs of corynoline and acetylcorynoline respectively.

**Figure S9.**
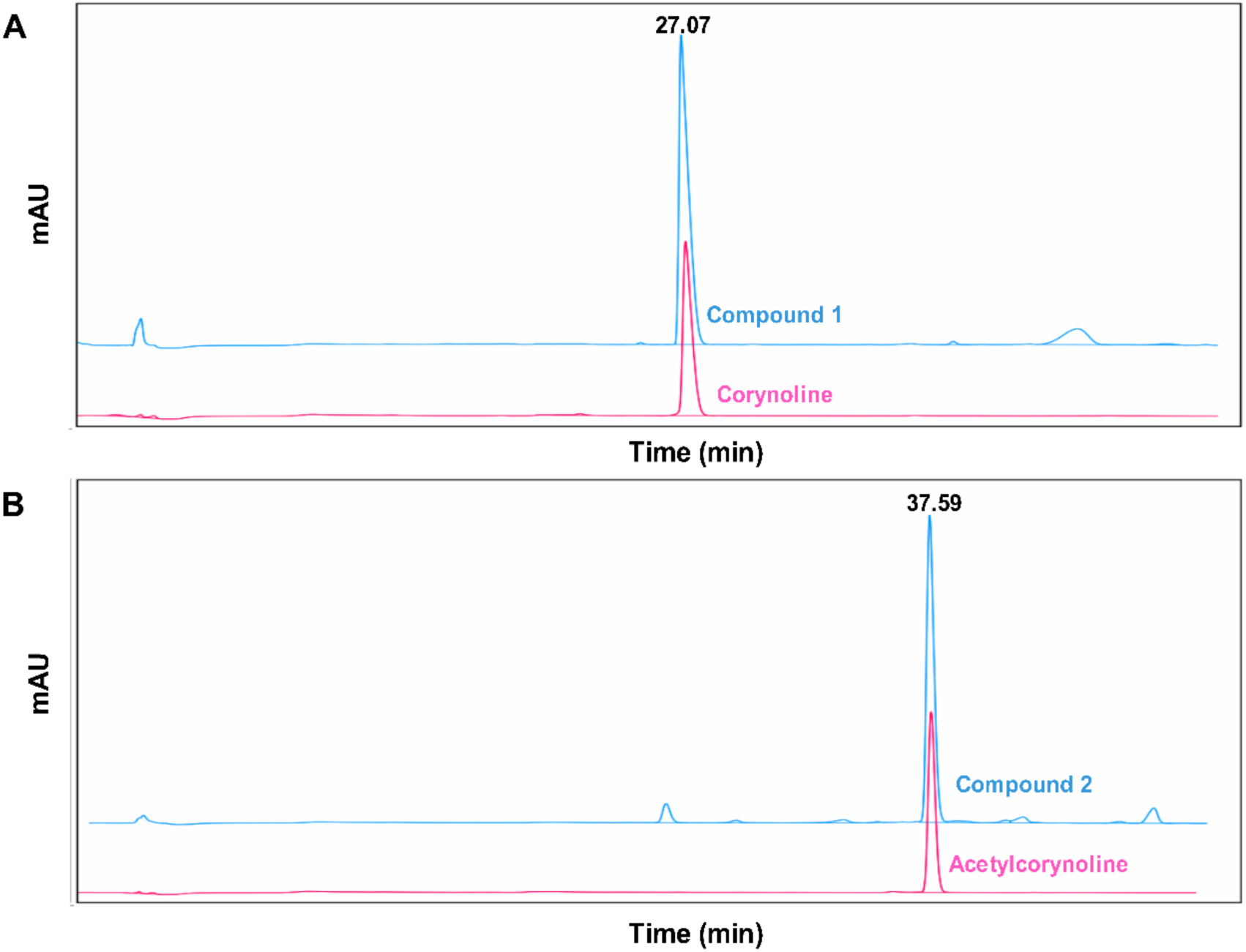
HPLC analysis of compounds 1 and 2, corynoline and acetylcorynoline. Purified natural compounds 1 and 2 together with commercially obtained corynoline and acetylcorynoline were analyzed by analytic HPLC. Both compound 1 and corynoline were eluted at 27.07 min (A), whereas compound 2 and acetylcorynoline were eluted at 37.59 min (B). Detection was performed at 280 nm.

**Figure S10.**
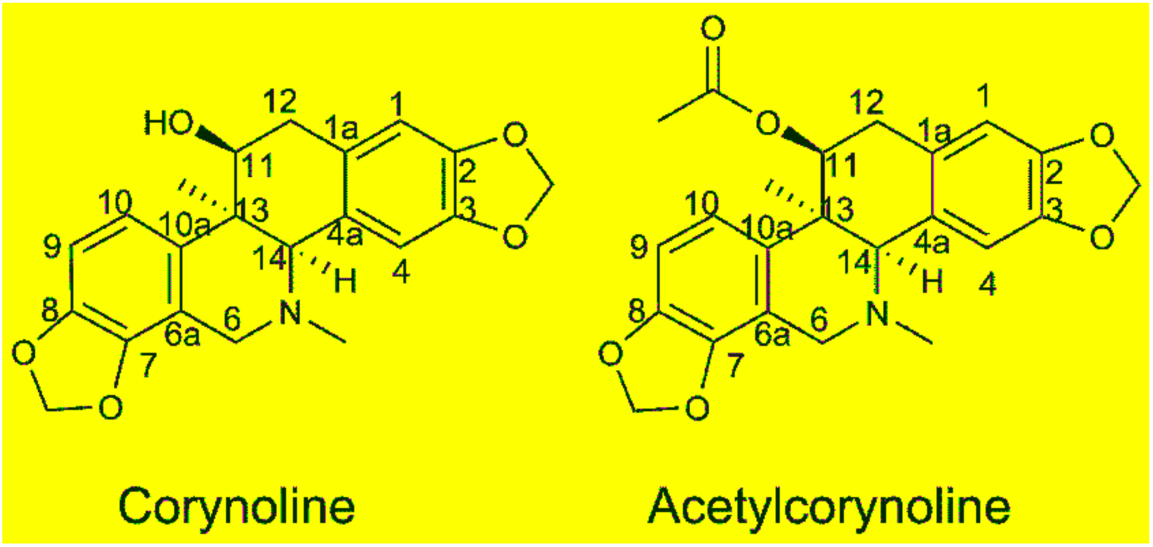
Chemical structures of corynoline and acetylcorynoline.

**Figure S11.**
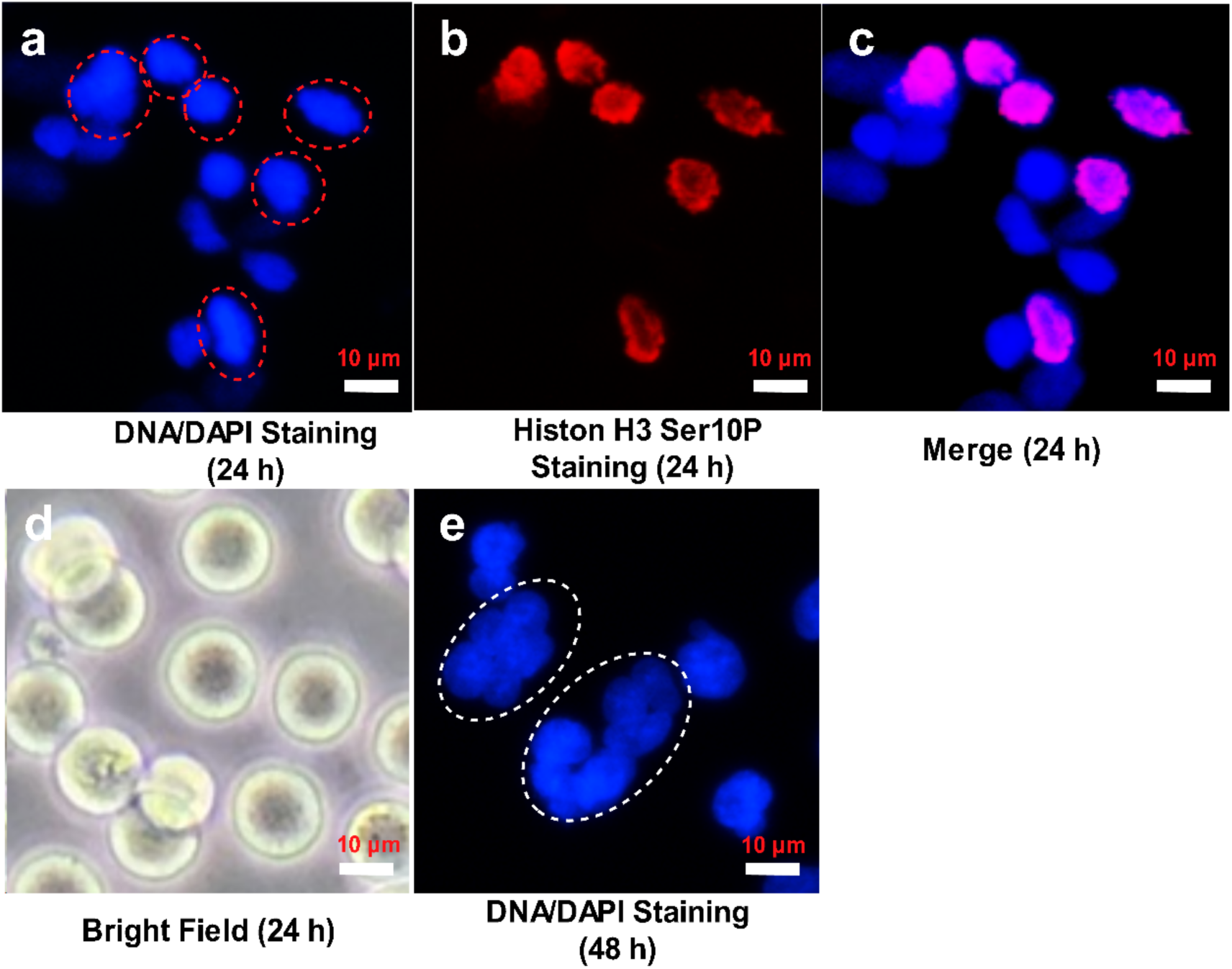
Corynoline arrests cells in early mitosis and elicits polyploidy. RPE-MBC cells were treated with 5 μM of commercially obtained corynoline for 24 hours (a-d) or 48 hours (e) before micrographed using an inverted microscope (d), or fixed and stained for DNA with DAPI (blue, a-c, e) or Histone H3 phosphorylated at Ser10 (red, b, c). Red circles denote mitotic cells with condensed nuclei, whereas white circles denote polyploid cells. Scale bars, 10 μm.

**Figure S12.**
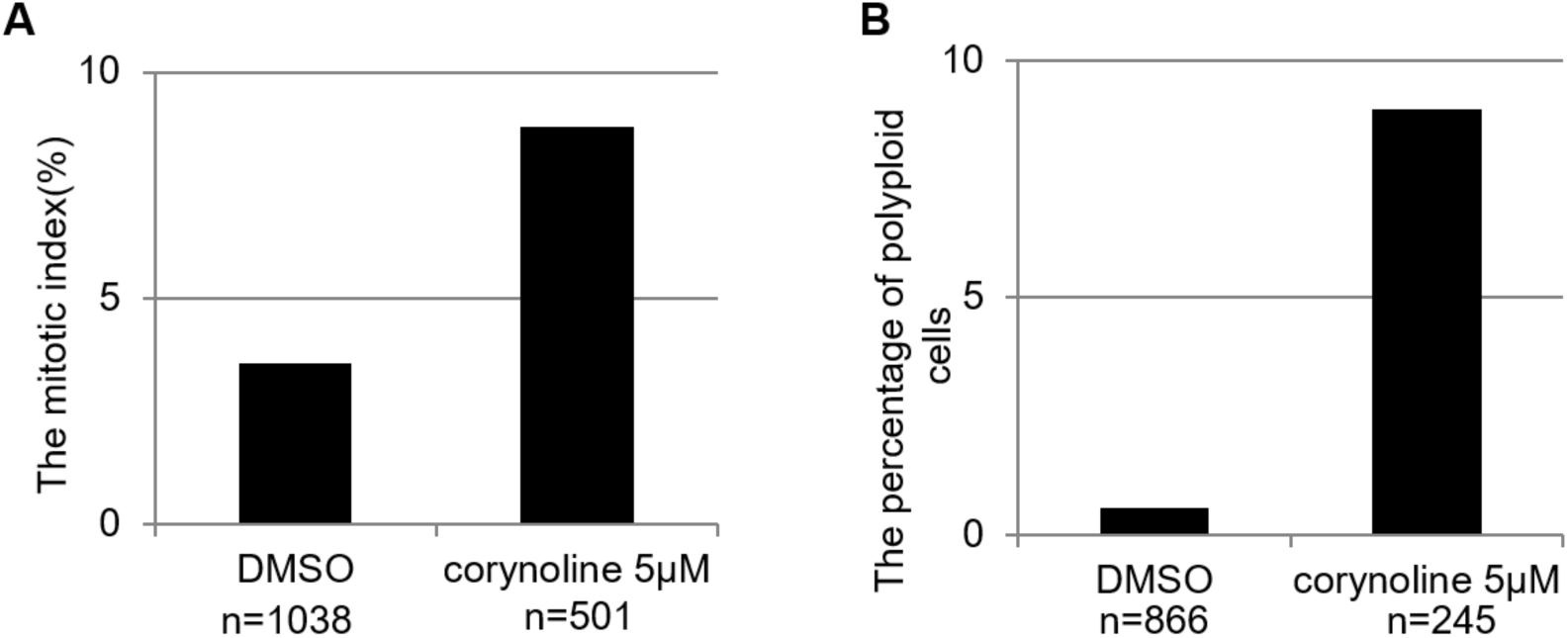
Quantitation of cells that were positive for Histone H3 phosphorylated at Ser10 at 24 hours and subsequent polyploidy at 48 hours. RPE-MBC cells were treated with 5 μM corynoline for 24 hours (A) or 48 hours (B) before fixation and staining. Graphs represent a single representative experiment where phosphorylation of Histone H3 at Ser10 serves to mark cells paused in mitosis (A) and polyploidy (B) was quantified in a paired culture one day later. In this instance, n indicates the number of cells observed per field of view.

**Figure S13.**
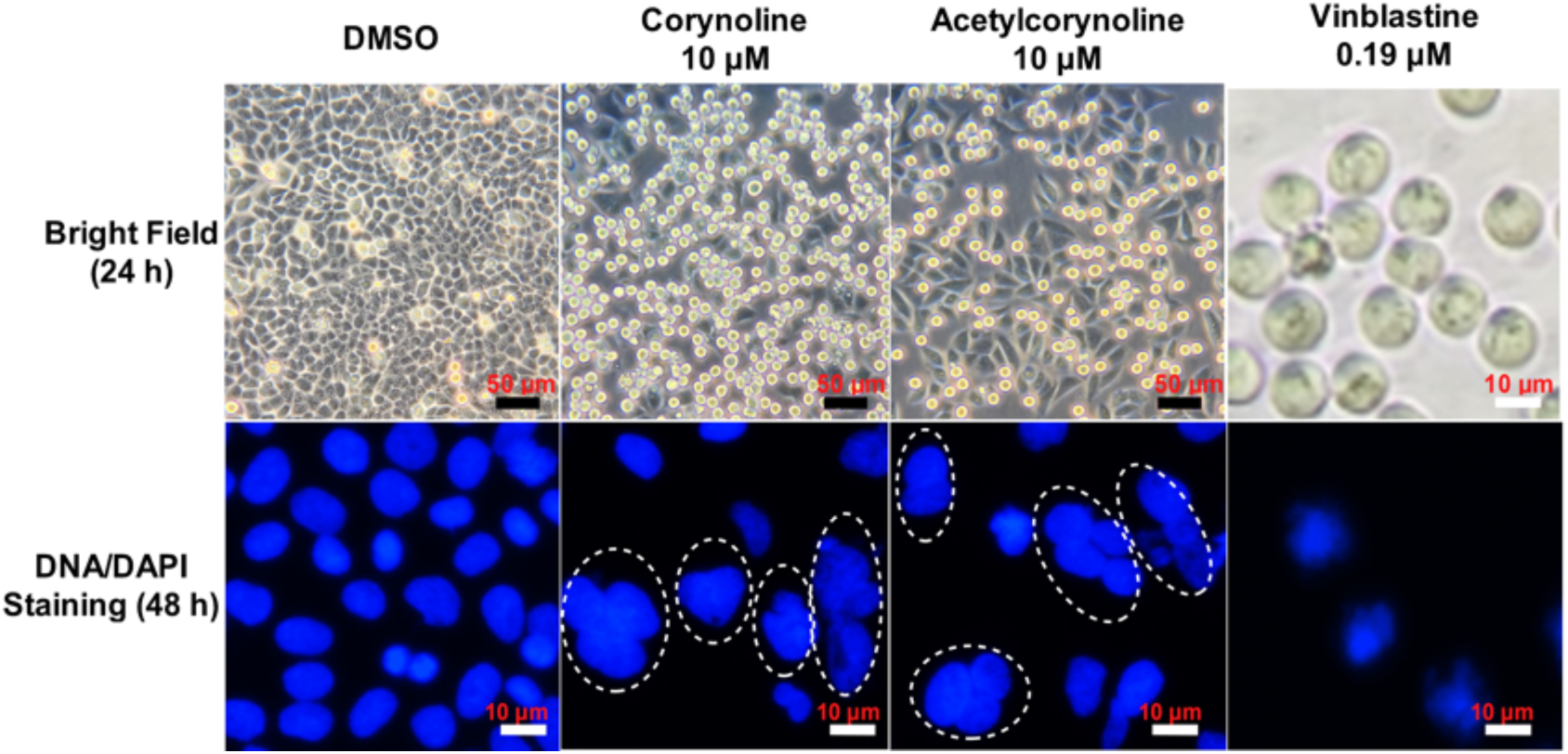
Both corynoline and acetylcorynoline provoke the spindle checkpoint response and elicit polyploidy. RPE-MBC cells were treated with the indicated concentrations of corynoline, acetylcorynoline or vinblastine (DMSO control) before visualization under an inverted tissue culture microscope at 24 hours or stained for DNA with DAPI at 48 hours. Polyploid cells are highlighted with white ovals.

**Table S1.** ^**1**^**H NMR (400 MHz) and** ^**13**^**C NMR (100 MHz) Data for compounds 1 and 2 in CDCl**_**3**_.

| No. | 1 |  | 2 |  |
| --- | --- | --- | --- | --- |
|  | δC | δH (*J* in Hz) | δC | δH (*J* in Hz) |
| 1 | 107.0 | 6.65, s | 108.2 | 6.52, s |
| 1a | 125.4 |  | 127.0 |  |
| 2 | 144.2 |  | 145.7 |  |
| 3 | 144.7 |  | 146.2 |  |
| 4 | 112.4 | 6.66, s | 112.4 | 6.88, s |
| 4a | 127.4 |  | 129.6 |  |
| 6 | 52.7 | 4.05, d (15.3)<br>3.45, d (15.3) | 48.15 | 3.91, d (15.3)<br>3.53, d (15.3) |
| 6a | 116.7 |  | 117.0 |  |
| 7 | 142.0 |  | 142.5 |  |
| 8 | 147.2 |  | 144.2 |  |
| 9 | 118.8 | 6.79, d (8.3) | 117.0 | 6.68, d (8.3) |
| 10 | 108.8 | 6.91, d (8.2) | 108.6 | 6.97, d (8.3) |
| 10a | 136.1 |  | 132.8 |  |
| 11 | 68.2 | 4.04, d | 68.2 | 5.22, t |
| 12 | 36.2 | 3.15, d<br>3.10, d | 32.0 | 2.97, d<br>2.86, d |
| 13 | 40.2 |  | 41.7 |  |
| 14 | 74.7 | 3.96, d | 74.6 | 3.51, d |
| N-CH<sub>3</sub> | 42.4 | 2.23, s | 43.1 | 2.48 s |
| 13-CH<sub>3</sub> | 23.3 | 1.15, s | 27.5 | 1.27, s |
| OCH<sub>2</sub>O | 100.8 | 5.95 m | 100.7 | 5.93, m |
| OCH<sub>2</sub>O | 101.0 | 6.00 m | 100.9 | 5.93, m |
| 11-OH/11-OAc |  |  | 169.7; 20.8 | 1.86, s |

**Table S2.** Corynoline and acetylcorynoline elicit mitotic arrest and induce polyploidy in a concentration-dependent manner.

| Concentration<br>(μM) | Corynoline | | Acetylcorynoline | | Taxol | | DMSO |
| --- | --- | --- | --- | --- | --- | --- | --- |
|  | R | P | R | P | R | P | R/P |
| 40 | + | + | + | + | + | - | - |
| 20 | + | + | + | + | + | - | - |
| 10 | + | + | + | + | + | - | - |
| 5 | + | + | - | - | + | - | - |
| 2.5 | - | - | - | - | + | - | - |
| 1.26 | - | - | - | - | + | - | - |
| 0.625 | - | - | - | - | + | - | - |
| 0.3125 | - | - | - | - | + | - | - |

R and P represents ‘round up’ and ‘polyploidy’ respectively. “+” and “-” indicate positive and negative, respectively, for the indicated phenotypes. Cell round-up was assayed at 24 hours and polyploidy at 48 hours after the initiation of treatment.

